# Evaluating cell lines as models for metastatic cancer through integrative analysis of open genomic data

**DOI:** 10.1101/337287

**Authors:** Ke Liu, Patrick A. Newbury, Benjamin S. Glicksberg, William ZD Zeng, Eran R. Andrechek, Bin Chen

**Affiliations:** Department of Pediatrics and Human Development, College of Human Medicine, Michigan State University, Grand Rapids, MI, USA; Department of Pharmacology and Toxicology, College of Human Medicine, Michigan State University, Grand Rapids, MI, USA; Bakar Computational Health Sciences Institute, University of California San Francisco, San Francisco, CA, USA; Department of Physiology, Michigan State University, East Lansing, MI, USA

## Abstract

Metastasis is the most common cause of cancer-related death and, as such, there is an urgent need to discover new therapies to treat metastasized cancers. Cancer cell lines are widely-used models to study cancer biology and test drug candidates. However, it is still unknown to what extent they adequately resemble the disease in patients. The recent accumulation of large-scale genomic data in cell lines, mouse models, and patient tissue samples provides an unprecedented opportunity to evaluate the suitability of cell lines for metastatic cancer research. In this work, we used breast cancer as a case study. The comprehensive comparison of the genetic profiles of 57 breast cancer cell lines with those of metastatic breast cancer samples revealed substantial genetic differences. In addition, we identified cell lines that more closely resemble different subtypes of metastatic breast cancer. Surprisingly, a combined analysis of mutation, copy number variation and gene expression data suggested that MDA-MB-231, the most commonly used triple negative cell line for metastatic breast cancer research, had little genomic similarity with Basal-like metastatic breast cancer samples. We further compared cell lines with organoids, a new type of preclinical model which are becoming more popular in recent years. We found that organoids outperformed cell lines in resembling the transcriptome of metastatic breast cancer samples. However, additional differential expression analysis suggested that both types of models could not mimic the effects of tumor microenvironment and meanwhile had their own bias towards modeling specific biological processes. Our work provides a guide of cell line selection in metastasis-related study and sheds light on the potential of organoids in translational research.

## Introduction

Cancer cell lines were initially derived from tumors and cultured in a 2D environment. Due to the merit of cell culture, they have been widely used as models to study cancer biology and test drug candidates^1^. However, the fact that many drugs with promising preclinical evidence fail in the clinic urges the reinvestigation of cell lines as tumor models^2^. The differences between cell lines and tumors have raised the critical question to what extent cell lines recapitulate the biology of tumor samples^3,4^.

The emergence of large-scale genomic data provides an unprecedented opportunity to quantify the biological differences between cancer cell lines and human tumors. The Cancer Genome Atlas (TCGA) project characterized both genetic and transcriptome profiles of more than 10,000 human tumor samples across over 32 tumor types^5^. The Cancer Cell Line Encyclopedia (CCLE) characterized both genetic and transcriptome profiles for more than 1,000 cell lines^6^. Silvia *et al*. performed comprehensive comparison of molecular profiles between 47 ovarian cancer cell lines and ovarian tumor samples and showed that none of popular cell line models closely resemble high-grade serous ovarian tumor samples^7^. In addition, they identified several rarely used cell lines closely resembled ovarian tumor samples. We examined the transcriptome similarity between hepatocellular carcinoma (HCC) cell lines and HCC tumor samples and demonstrated that nearly half of the HCC cell lines did not resemble HCC tumor samples^8^. Jian *et al*. conducted a comprehensive comparison of molecular portraits between breast cancer cell lines and primary breast cancer samples, and uncovered both similar and dissimilar molecular features^9^.

Cancer metastasis is the most common cause of cancer-related death, thus there is an urgent need of new drugs for treating cancer metastasis^10,11^. Previous cell line evaluation analysis was mainly performed in reference to primary tumors. It remains unknown whether cell lines closely resemble metastatic cancer and thus are appropriately used in translational research. Robinson *et al*. performed whole-exome and transcriptome sequencing on about 500 metastatic cancer samples and recently released their dataset (refer to MET500)^12^. This large-scale genomic profiling combined with existing genomic data allows the evaluation of the suitability of cell lines as models for metastatic cancer. Using breast cancer as a case study, we comprehensively compared multiple types of molecular features between breast cancer cell lines and metastatic breast cancer samples (Fig 1). Based on our analyses, we identified cell lines that are suitable for modeling metastatic breast cancer samples of individual subtypes. In addition, we evaluated patient-derived organoids and showed their potential in pre-clinical studies. Our work provides useful guidance for choosing cell lines in metastasis-related translational research and could be easily extended to other cancer types.

**Fig 1.**
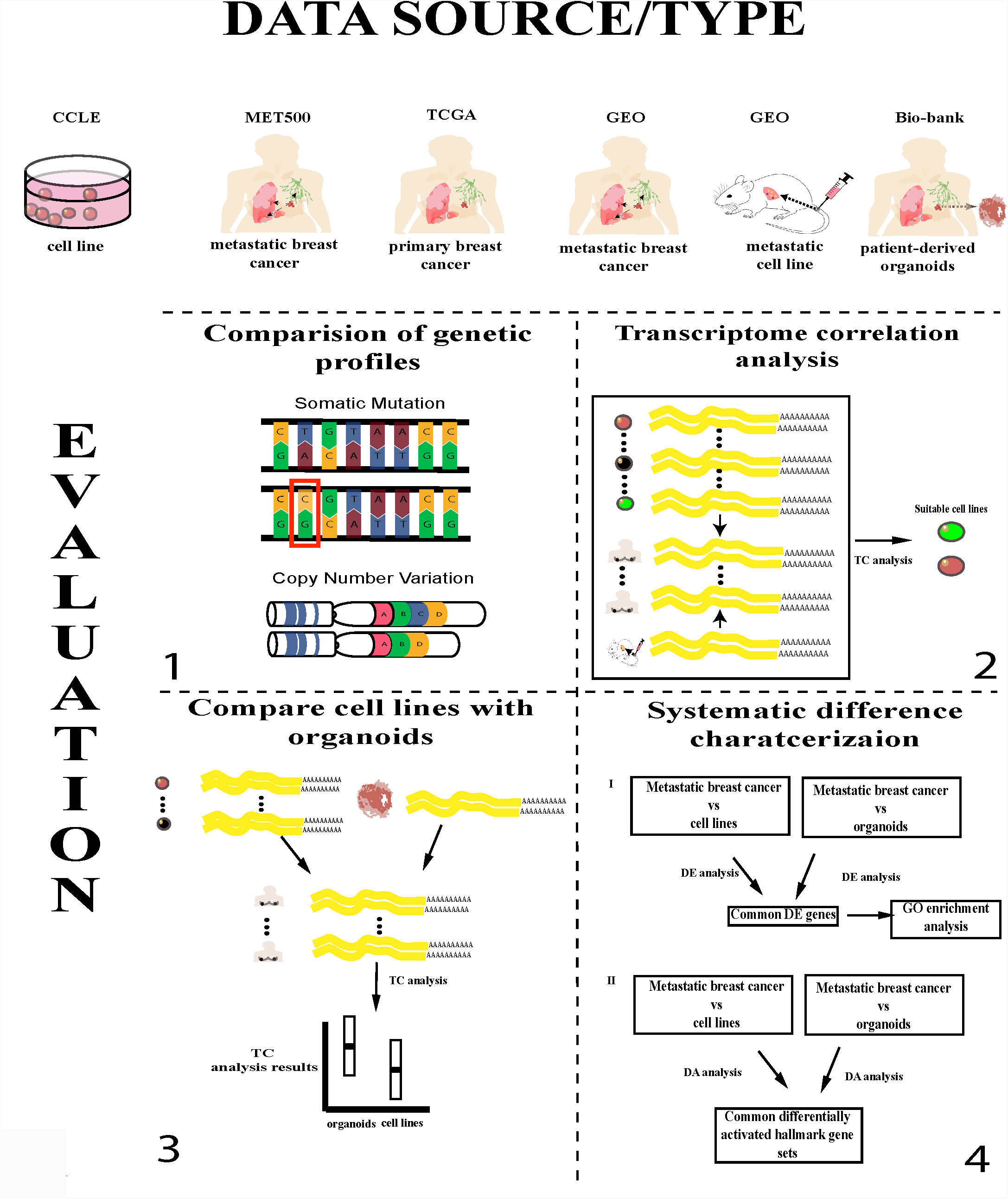
Overall design of the study. The upper panel lists data sources as well as sample types used in our study and the lower panel is a summary of the evaluations performed. TC analysis: transcriptome correlation analysis; DE analysis: differential gene expression analysis; DA analysis: gene set differential activity analysis.

## Results

### Comparison of genetic profiles between metastatic breast cancer and cell lines

We first compared somatic mutation profiles between MET500 breast cancer samples and CCLE breast cancer cell lines. Whole-exome sequencing was performed for MET500 samples, while hybrid capture sequencing was performed for CCLE cell lines. We thus only focused on the 1,630 genes genotyped in both studies. We were particularly interested in two types of genes that may play important roles in breast cancer metastasis: genes that are highly mutated in metastatic breast cancer, and genes that are differentially mutated between metastatic and primary breast cancers.

Consistent with previous research, we identified a long-tailed mutation spectrum of the 1,630 genes in MET500 breast cancer samples and 69 of them were highly mutated (mutation frequency > 0.05, Fig S1a). The five most-altered genes were TP53 (0.67), PIK3CA (0.35), TTN (0.29), OBSCN (0.19), and ESR1 (0.14). We identified 19 differentially mutated genes between MET500 and TCGA samples (FDR < 0.001) and the five most significant genes were ESR1, TNK2, OBSN, CAMKK2, and CLK1 (Fig S1b). Interestingly, all of these 19 differentially mutated genes had higher mutation frequency in MET500 than TCGA, which is consistent with previous study showing that metastatic cancer has increased mutation burden compared to primary cancer^12^. 68% of them were also among the 69 highly mutated genes mentioned above. After merging the two gene lists, 75 unique genes remained (Fig 2a and Table S1). The median mutation frequency of the 75 genes across CCLE breast cancer cell lines was 0.07 and only 9% of them (PRKDC, MAP3K1, TTN, ADGRG4, TP53, FN1, and AKAP9) were mutated in at least 50% of cell lines, suggesting that majority of these gene mutations could be recapitulated by only a few cell lines. Surprisingly, nine of the 75 genes (ESR1, GNAS, PIKFYVE, FFAR2, RNF213, MYBL2, KAT6A, MAP4K4, and FMO4) were not mutated in any cell line. Notably, ESR1 has been identified as a driver gene of cancer metastasis and associated mutations can cause endocrine resistance of metastatic breast cancer cells^13,14^, yet none of the cell lines could be used to appropriately model it.

**Fig 2.**
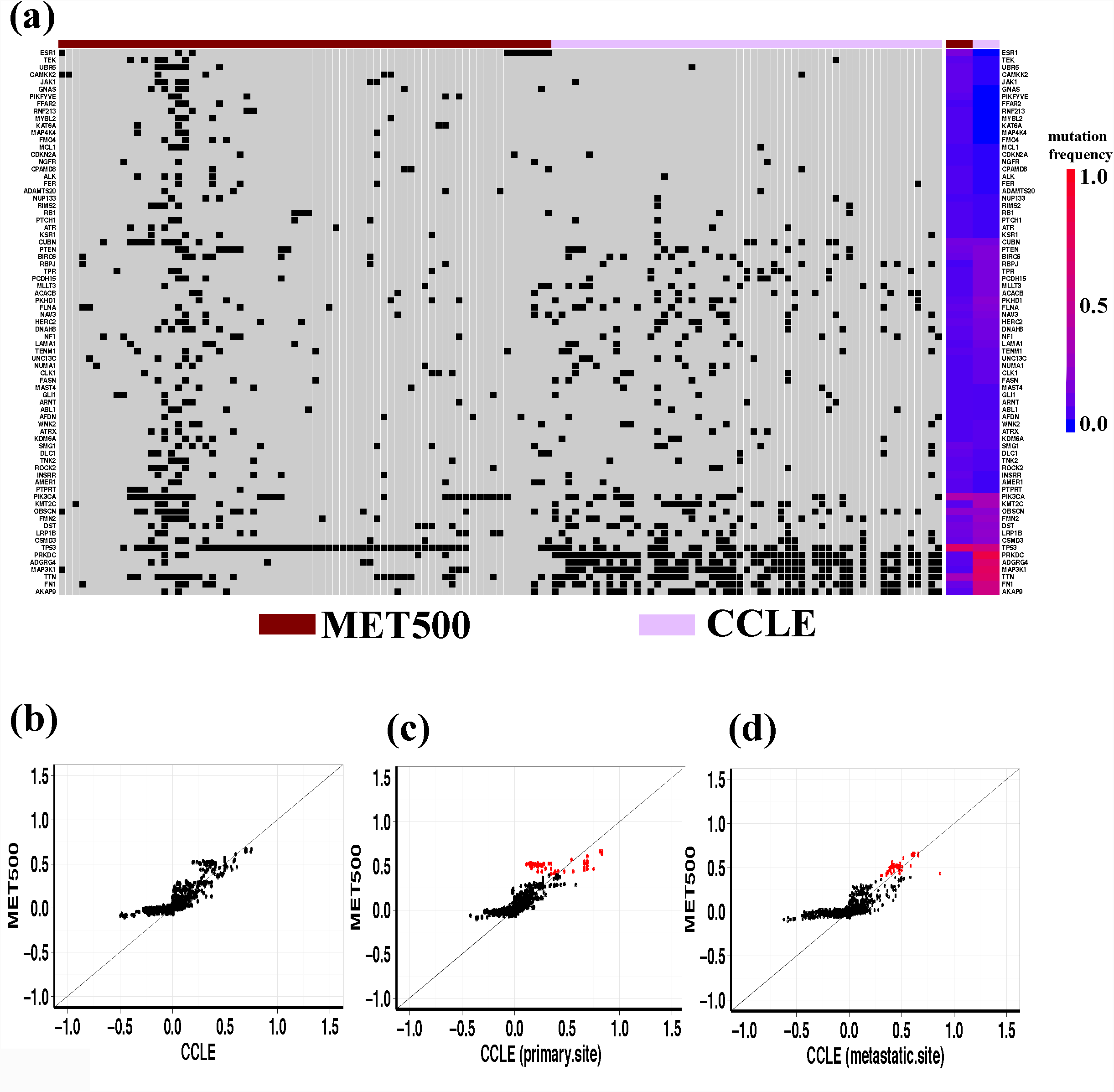
Comparison of genetic profiles between MET500 breast cancer samples and CCLE breast cancer cell lines. (a) Somatic mutation profile of the 75 genes across MET500 breast cancer samples and CCLE breast cancer cell lines. The top-side color-bar indicates data sources (MET500 or CCLE) and the right-side color-bar indicates mutation frequency. (b) Comparison of CNV profiles between MET500 breast cancer samples and CCLE breast cancer cell lines. (c) Comparison of CNV profiles between MET500 breast cancer samples and the 33 primary-site derived CCLE breast cancer cell lines. (d) Comparison of CNV profiles between MET500 breast cancer samples and the 24 metastatic-site derived CCLE breast cancer cell lines. In panel (b), (c) and (d), each dot is a gene, with y-axis representing its median CNV value across MET500 breast cancer samples, and x-axis representing its median CNV value across CCLE breast cancer cell lines. In panel (c) and (d), genes with high copy-number-gain in MET500 breast cancer samples were marked by red.

We next asked whether there were genes specifically hypermutated in breast cancer cell lines. To address this question, we examined the mutation spectrum of the 32 genes that were mutated in at least 50% of the breast cancer cell lines. Surprisingly, 25 of them had low mutation frequency (< 0.05) in MET500 breast cancer samples. Further analysis of somatic mutation profiles of the 25 genes in TCGA breast cancer samples confirmed their hypermutations were specific to breast cancer cell lines (Fig S1c).

In addition to somatic mutation spectrum, we also compared copy number variation (CNV) profiles between MET500 breast cancer samples and CCLE breast cancer cell lines. We observed a high correlation of median-CNV values across the 1,630 commonly genotyped genes (spearman rank correlation = 0.81, Fig 2b). However, we also noticed that the gain-of-copy-number events in cell lines appeared to resemble metastatic breast cancer while loss-of-copy-number events did not. For the 711 genes showing copy-number-loss in CCLE breast cancer cell lines (median-CNV < 0), their cell line derived median-CNV values were significantly lower than that from MET500 breast cancer samples; however, no significant difference was detected in the 919 genes with copy-number-gain (Fig S1d).

Out of the 57 CCLE breast cancer cell lines, 24 were derived from metastatic sites (Table S2). We further divided the cell lines into two groups (according to whether they were derived from metastatic sites or not) and then compared the CNV profile of each group with MET500 breast cancer samples. We found cell lines derived from metastatic sites more closely resembled the CNV status of the 109 genes with high copy-number-gain (median-CNV >= 0.4) in MET500 breast cancer samples (Fig 2c, 2d, and Fig S1e).

### Correlating CCLE breast cancer cell lines with metastatic breast cancer samples using transcriptome data

Transcriptome correlation analysis (TC analysis) is proven to be an effective approach to evaluate the suitability of cell lines for research purpose^7,8,15^. Therefore, we performed TC analysis and ranked all 1,019 CCLE cell lines according to their transcriptome-similarly with MET500 breast cancer samples (see Methods). The top 20 cell lines were all breast cancer cell lines, suggesting metastatic breast cancer cells retain a transcriptomic signature from the tissue they originated in and cell lines have the potential to resemble the transcriptome of them (Fig 3a). MDA-MB-415 and HMC18 were the two breast cancer cell lines that had highest and lowest transcriptome-similarity, respectively (0.415 and 0.087).

**Fig 3.**
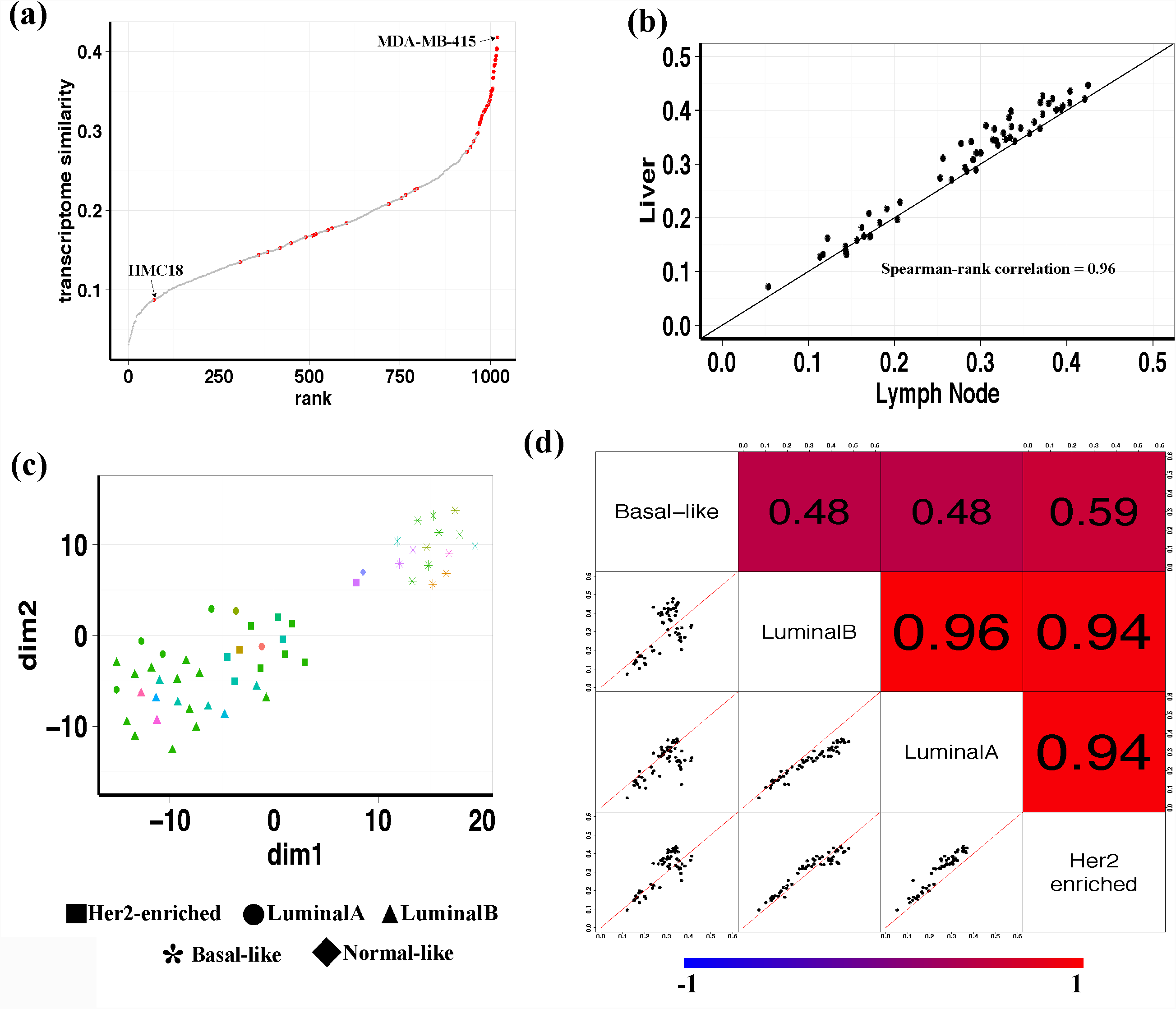
TC analysis between MET500 breast cancer samples and CCLE breast cancer cell lines. (a) 1,019 CCLE cell lines are ranked according to their transcriptome-similarity with MET500 breast cancer samples. Each dot is a CCLE cell line and breast cancer cell lines are marked by red. (b) Metastatic-site-specific TC analysis results are highly correlated between liver and lymph node. Each dot is a CCLE breast cancer cell line, with x-axis representing its transcriptome-similarity with the 9 lymph node derived MET500 breast cancer samples, and y-axis representing its transcriptome-similarity with the 27 liver derived MET500 breast cancer samples. (c) t-SNE plot of MET500 breast cancer samples. Metastatic-sites are labeled by color and subtypes are labeled by shape. (d) Pair-wise comparison of subtype-specific TC analysis results. In the lower-left plots, each dot is a CCLE breast cancer cell line, with the two axis representing transcriptome-similarity of the cell line with MET500 breast cancer samples of the two intersecting subtypes. The upper-right shaded values are the corresponding pair-wise spearman rank correlation values of each pair.

We next assessed whether cell lines resembling the transcriptome of samples from different metastatic sites were identical. We were only able to consider liver and lymph node (the two sites which have at least nine samples) due to the lack of enough samples from other sites in the MET500 dataset. For each of them, we performed metastatic-site-specific TC analysis (i.e., compute transcriptome-similarity of cell lines with samples derived from a specific metastatic site) and found the results were highly correlated (Fig 3b) with MDA-MB-415 being the most correlated cell line for both sites. In addition, we detected no significant difference of expression correlation (with MDA-MB-415) between the two sites (Fig S2a).

Given the genomic heterogeneity of breast cancer, we further asked whether cell lines resembling the transcriptome of metastatic breast cancer of different subtypes were identical. To address this question, we first determined the PAM50 subtype of MET500 breast cancer samples with R package genefu then applied t-SNE to visualize them (Fig 3c). We found Basal-like samples clustered together and separated from other subtypes; additionally, the majority of LuminalA/LuminalB/Her2-enriched/Normal-like samples were mixed together except two skin-derived samples. HER2-enriched samples seemed to be separated from LuminalA/LuminalB samples but the boundary was not clear. These results suggested that subtype information was well maintained in metastatic breast cancer samples and additionally confirmed the feasibility of using PAM50 for subtyping metastatic breast cancer though it was initially developed using primary breast cancer data. We further confirmed the subtyping results by performing the same analysis on a combined dataset which contains both MET500 and TCGA breast cancer samples (Fig S2b). Next, we performed subtype-specific TC analysis (i.e., compute transcriptome-similarity of cell lines with samples of a specific subtype) and found high correlation within LuminalA/LuminalB/Her2-enriched subtypes, in contrast to their relatively lower correlation to Basal-like subtype (Fig 3d).

To confirm the robustness of our TC analysis on MET500 dataset, we searched the GEO database and assembled a microarray dataset containing the expression value of another 103 metastatic breast cancer samples, and then repeated the above analysis. Results obtained from two the datasets were highly consistent with each other. First, there was a large overlap of the top-ranked cell lines. Out of the 10 cell lines having highest transcriptome-similarity with the 103 metastatic breast cancer samples, six of them were within the 10 cell lines having highest transcriptome-similarity with MET500 breast cancer samples. Second, both metastatic-site-specific and subtype-specific TC analysis results showed high correlations (Fig S3). Due to such high consistency, it is not surprising that we observed similar correlation trends in metastatic-site-specific (and subtype-specific) TC analysis results (Fig S4, S5).

About 24% of the 103 samples in the microarray dataset were derived from bone. Remarkably, the metastatic-site-specific TC analysis result of bone showed lower correlation with other sites (Fig S4). To exclude the possibility that this was caused by tumor purity issues, we applied ESTIMATE^16^ on the microarray dataset and found the tumor purity of bone-derived samples was not significantly lower than that of liver, lymph node, and lung (Fig S6). Our results may not be too surprising given the fact that bone provides a very unique microenvironment including enriched expression of osteolytic genes^17^; however, this result needs to be confirmed in the future as more data becomes available.

### Suitable cell lines for metastatic breast cancer research

We attempted to identify suitable cell lines for different subtypes of metastatic breast cancer based on the results of subtype-specific TC analysis. Given a subtype, we noticed that for a random CCLE cell line, its transcriptome-similarity with MET500 breast cancer samples of that subtype approximately followed a normal distribution (Fig S7). Therefore, those breast cancer cell lines showing significantly higher transcriptome-similarity were considered as suitable. Driven by this finding, for each subtype we first fit a normal distribution (the null distribution) with the transcriptome-similarity values (derived from subtype-specific TC analysis) of all non-breast-cancer cell lines and then assigned each of the 57 breast cancer cell lines a right-tailed p-value. The most significant cell lines for LuminalA, LuminalB, Her2-enriched, and Basal-like subtypes were MDA-MB-415 (p-value =3.59e-05), BT483 (p-value=2.22e-07), EFM192A (p-value=0.11e-and HCC70 (p-value =0.40e-03), respectively. Using a criteria of FDR <= 0.01, we identified 20, 28 and 24 suitable cell lines for LuminalA, LuminalB, and Her2-enriched subtypes respectively. Notably, most of these significant cell lines were derived from metastatic sites and 18 were shared by the three subtypes. Surprisingly, no cell line passed the criterion for Basal-like subtype. We further examined whether this was due to the limited number of Basal-like MET500 breast cancer samples, but found that the number of LuminalA samples was even less than that of Basal-like samples. After we used a more loosened FDR cutoff of 0.05, we obtained 22 suitable cell lines for Basal-like subtype. All statistical testing results are listed in Table S3.

We next searched PubMed to examine the popularity of the 57 breast cancer cell lines (see Methods and Table S2). MCF7 is most commonly used in metastatic breast cancer research (43.6% of total PubMed citations). Although we found it was a suitable cell line for LuminalB subtype, it was less correlated with LuminalB MET500 breast cancer samples than BT483 (Fig S8a). Following MCF7 is MDA-MB-231 (40.2% of total PubMed citations); however, it was not a suitable cell line for any subtype. The third most commonly used cell line was T47D (3.9% of total PubMed citations), which was a suitable cell line for LuminalA and Her2-enriched subtypes. It did not show significantly lower correlation with LuminalA MET500 breast cancer samples than MDA-MB-415 (Fig S8b); however, compared to EFM192A, it was significantly less correlated with Her2-enriched MET500 breast cancer samples (Fig S8c).

While the triple negative cell line MDA-MB-231 is one of the most frequently used cell lines in metastatic breast cancer research, it might not be the most suitable cell line to model metastasis biology in breast cancer. We ranked all of the 1,019 CCLE cell lines according to their transcriptome-similarity with Basal-like MET500 breast cancer samples and the rank of MDA-MB-231 was 583 (Fig 4a). Consistent with this, MDA-MB-231 was significantly less correlated with Basal-like MET500 breast cancer samples than HCC70. We observed similar patterns with CNV data (Fig 4b). We also examined how MDA-MB-231 recapitulated the somatic mutation spectrum of Basal-like metastatic breast cancer samples and found only three of the 25 highly-mutated genes (mutation frequency >= 0.1 in Basal-like MET500 breast cancer samples) were mutated in MDA-MB-231 (Fig 4c). Since CCLE data for MDA-MB-231 was generated *in vitro*, to confirm our finding we obtained another independent microarray dataset which profiled the gene expression of MDA-MB-231 cell lines derived from lung metastasis *in vivo*^18^. We found, however, that even these MDA-MB-231 cell lines *in vivo* did not most closely resemble the transcriptome of lung metastasis breast cancer samples. The breast cancer cell line which showed highest correlation with lung metastasis breast cancer samples was EFM192A (Fig 4d). Our analysis indicates that although MDA-MB-231 presents many favorable properties for metastatic breast cancer research, its genomic profile is substantially different from metastatic breast cancer samples.

**Fig 4.**
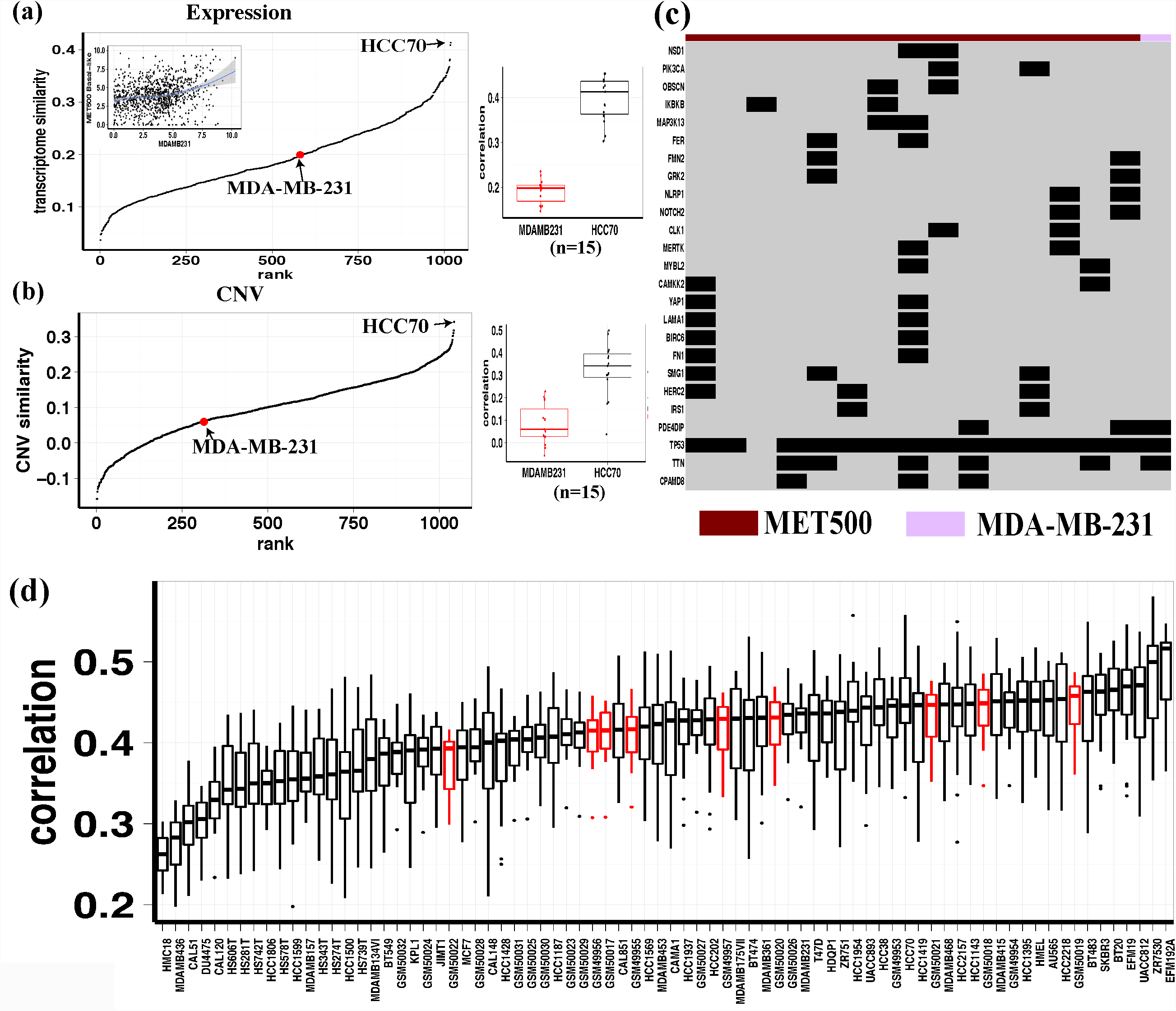
MDA-MB-231 has substantial genomic difference with Basal-like metastatic breast cancer samples. (a) The left panel shows the ranking of all 1,019 CCLE cell lines according to their transcriptome-similarity with Basal-like MET500 breast cancer samples. The top-left scatter plot within the first panel shows the expression of the most-varied 1,000 genes, with x-axis representing expression value in MDA-MB-231, and y-axis representing median expression value across Basal-like MET500 breast cancer samples. The boxplot on the right shows the distribution of expression correlation with Basal-like MET500 breast cancer samples for MDA-MB-231 and HCC70. (b) The left panel shows the ranking of all 1,019 CCLE cell lines according to their CNV similarity with Basal-like MET500 breast cancer samples; the boxplot on the right shows the distribution of CNV correlation with Basal-like MET500 breast cancer samples for MDA-MB-231 and HCC70. (c) Somatic mutation profile of the 25 highly mutated genes across MDA-MB-231 and Basal-like MET500 breast cancer samples. (d) Boxplot of expression correlation between cell lines and lung-derived metastatic breast cancer samples. This includes CCLE breast cancer cell lines and lung-metastasis-derived MDA-MB-231 (colored red).

### Recently established patient-derived organoids more closely resemble the transcriptome of metastatic breast cancer samples

Owing to the advancement of 3D culture technology, more and more tumor patient-derived organoids have been established and widely used in translational research^19,20^. However, their suitability to model metastatic cancer has not been comprehensively evaluated with large-scale genomic data. To fill this gap, we performed additional TC analysis on 26 patient-derived organoids using RNA-Seq data. The aforementioned subtype-specific TC analysis showed that the Basal-like subtype had relatively lower correlation with other subtypes and we also observed similar trend in organoids (Fig 5a). We next asked whether organoids outperformed cell lines in resembling the transcriptome of metastatic breast cancer. For each of the non-Basal-like organoids, we computed its transcriptome-similarity with non-Basal-like MET500 breast cancer samples and found organoids had significantly higher transcriptome-similarity than CCLE breast cancer cell lines (Fig 5b, left panel). The superiority of organoids was also observed in the TC analysis of Basal-like subtype (Fig 5b, right panel). The previous analysis revealed that MDA-MB-415, BT483 and EFM192A were the three most suitable cell lines for LuminalA, LuminalB and Her2-enriched subtypes, respectively. Interestingly, for all the three subtypes MMC01031 was the organoid showing highest transcriptome-similarity and had significantly higher correlation with MET500 breast cancer samples than the corresponding most suitable cell line. Organoid W1009 had the highest transcriptome-similarity with Basal-like MET500 breast cancer samples and the expression correlation values were also significantly higher than HCC70, a triple-negative cell line that is most suitable for Basal-like subtype (Fig 5c).

**Fig 5.**
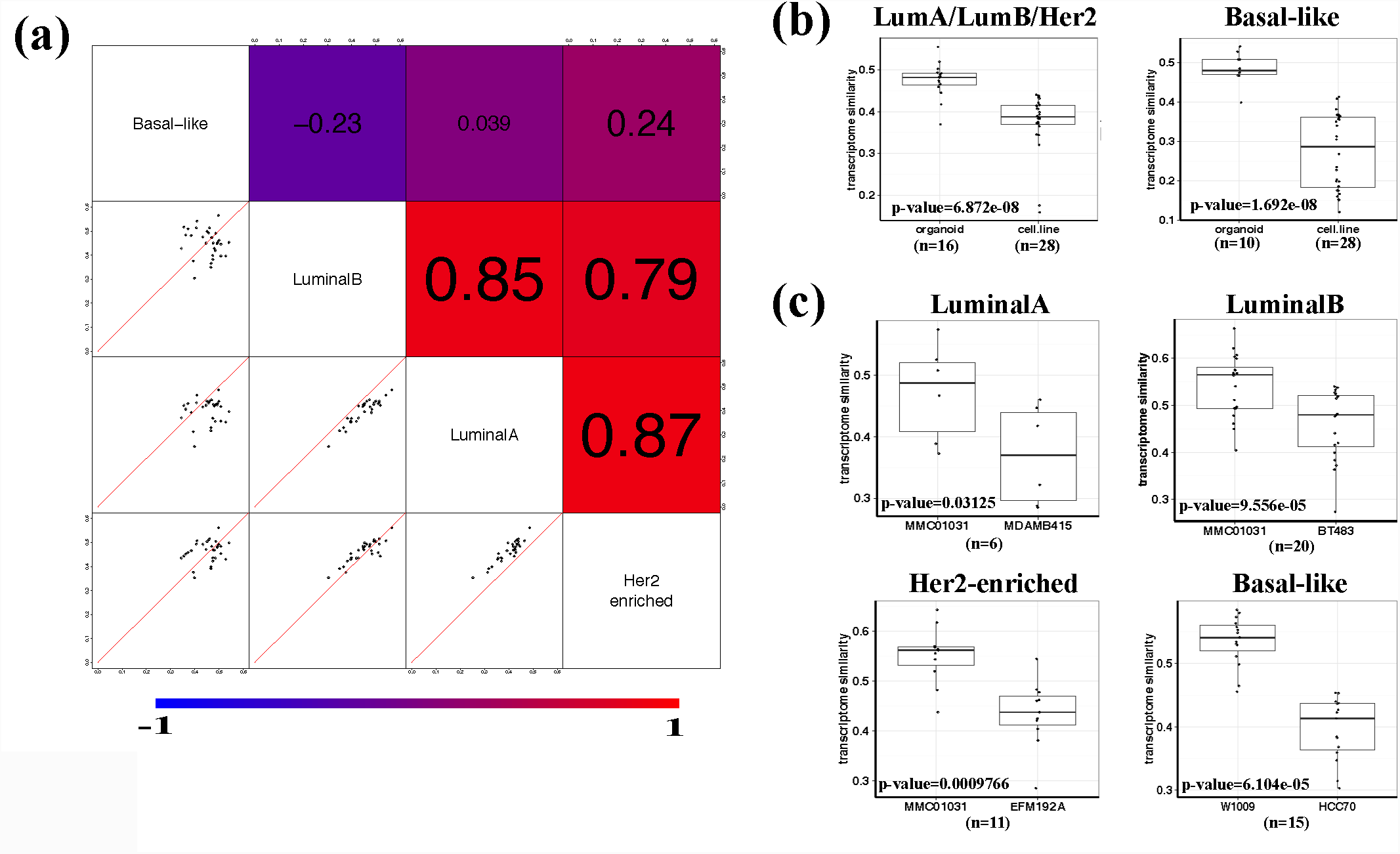
Comparing CCLE breast cancer cell lines with patient-derived organoids using gene expression data. (a) Pair-wise comparison of subtype-specific TC analysis results. In the lower-left plots, each dot is an established organoid, with the two axis representing transcriptome-similarity of the organoid with MET500 breast cancer samples of the two intersecting subtypes. The upper-right shaded values are the corresponding pair-wise spearman rank correlation values of each pair. (b) Boxplot of transcriptome-similarity (with MET500 breast cancer samples of different subtypes) of CCLE breast cancer cell lines and organoids. (c) For each subtype, the most-correlated organoid has significantly higher expression correlation with MET500 breast cancer samples of that subtype than the most-correlated cell line.

### Characterization of gene expression difference between metastatic breast cancer samples and *in vitro* models

Our TC analysis has shown that *in vitro* models such as cell lines and organoids could resemble the transcriptome of metastatic breast cancer at some extent. However, they are still different in many aspects. To characterize such differences, we performed differential gene expression analysis among MET500 breast cancer samples, CCLE breast cancer cell lines and organoids (Fig S9). For non-Basal-like subtypes, 2,380 genes (2,179 up-regulated, 201 down-regulated) were identified as differentially expressed in both MET500-vs-CCLE and MET500-vs-organoids comparisons. For Basal-like subtype, there were 1,378 common differential expressed (DE) genes (1,117 up-regulated, 261 down-regulated). After intersecting the above two common DE gene lists, we finally obtained 1,016 subtype-and-model-independent DE genes (948 up-regulated, 68 down-regulated) and then performed GO enrichment analysis. For the 948 up-regulated ones, 30 GO terms were identified as significant (FDR < 0.001) and most of them were immune-related, illustrating the large gap between culture media and tumor microenvironment (Table S4). The two terms “platelet degranulation”, and “chemotaxis” were also detected as significant. Besides microenvironment, our results also implicated the difference of intrinsic characteristics between metastatic breast cancer cells and *in vitro* models. For example, the enrichment on “steroid metabolic process” suggested that neither cell lines nor organoids resembled the reprogrammed metabolism of metastatic breast cancer sufficiently. Likewise, the enrichment on “cell adhesion” indicated that the *in vitro* models may not recapitulate epithelial-to-mesenchymal-transition-related process of metastatic breast cancer. Surprisingly, for the 68 down-regulated subtype-and-model-independent DE genes, no GO terms passed the FDR < 0.001 criteria, which could be due to the small gene number. We decreased the FDR cutoff to 0.1 and observed five significant terms with “cell division” being the most significant (FDR = 0.029).

We further performed gene set differential activity (DA) analysis on ssGSEA scores of the 50 MSigDB hallmark gene sets to characterize differences regarding to specific biological process (Fig 6a, Fig S10). For non-Basal-like subtype, we identified 35 and 32 significant gene sets in MET500-vs-CCLE and MET500-vs-organoids comparisons, respectively (FDR < 0.001, Table S5). There were 26 DA gene sets in common and for 23 of them the p-values derived from MET500-vs-CCLE comparison were lower than that derived from MET500-vs-organoid comparison, which may be unsurprising given that organoids more closely resemble the transcriptome of metastatic breast cancer samples (Fig 6b, left panel). We also performed DA analysis for Basal-like subtype, identifying 19 and 24 significant gene sets in MET500-vs-CCLE and MET500-vs-organoids comparisons, respectively (Fig 6b, right panel). For each of the subtypes, we classified the 50 hallmark gene sets into 4 categories according to DA analysis results:

**Fig 6.**
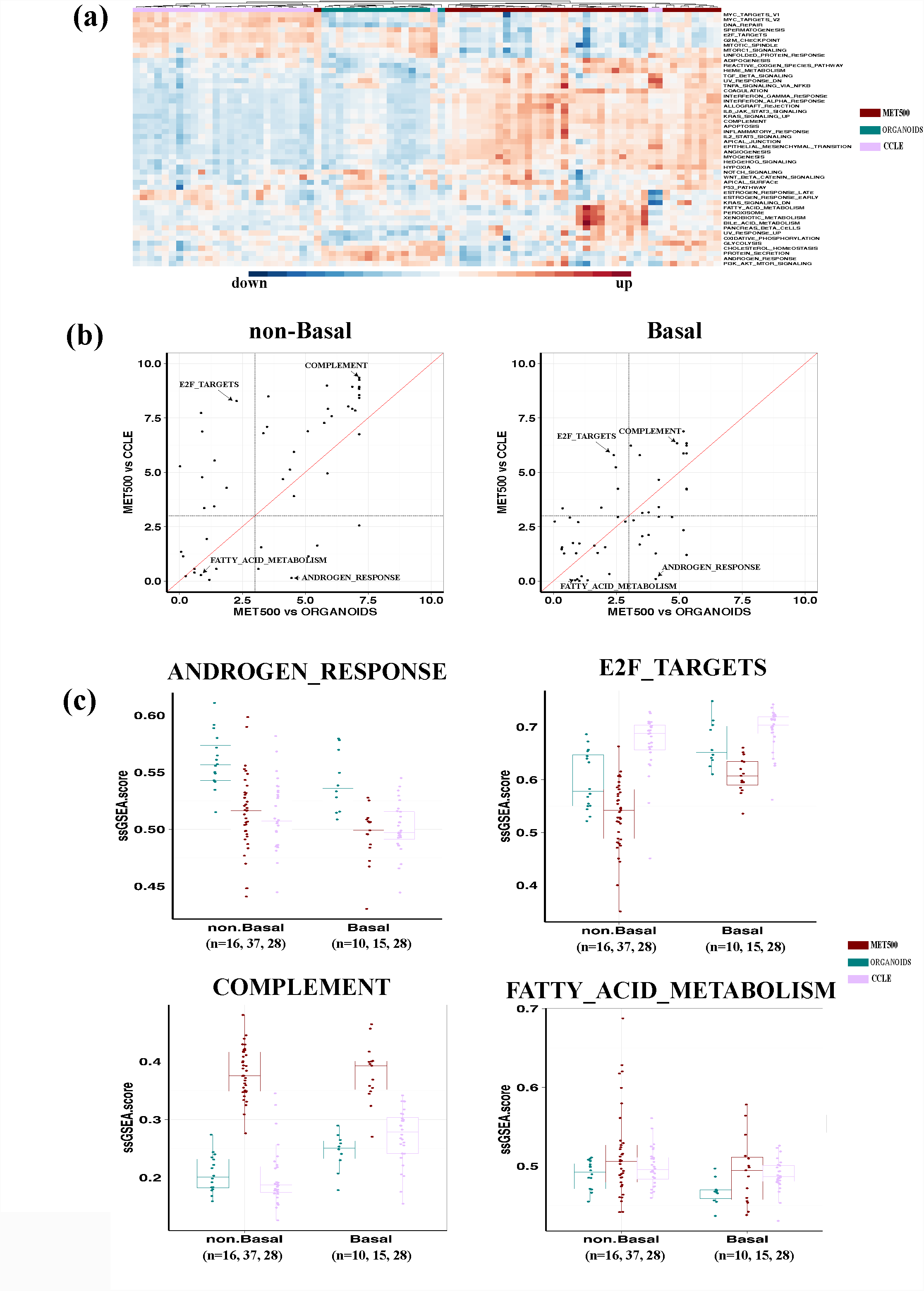
Comparison of ssGSEA scores of the 50 MSigDB hallmark gene sets. (a) Visualization of ssGSEA scores across non Basal-like CCLE breast cancer cell lines, MET500 breast cancer samples and organoids. (b) DA analysis results of different breast cancer subtypes. Each dot is a hallmark gene set, with x-axis representing −log10(FDR) derived from MET500-vs-organoids comparison, and y-axis representing −log10(FDR) derived from MET500-vs-CCLE comparison. (c) Boxplot of ssGSEA scores of the four representative gene sets.

I. Only significant in MET500-vs-organoids comparison (e.g., ANDROGEN RESPONSE).
II. Only significant in MET500-vs-CCLE comparison (e.g., E2F TARGETS).
III. Significant in both MET500-vs-organoids and MET500-vs-CCLE comparisons (e.g, COMPLEMENT).
IV. Not significant in either comparison (e.g, FATTY ACID METABOLISIM).

Interestingly, 27 gene sets could be consensually classified into one specific category, regardless of the subtype. Fig 6c shows the distribution of ssGSEA scores of the representative gene set for each category.

## Discussion

In cancer research, cell lines have been traditionally used to test drug candidates and study disease mechanism. The genetic profile comparison showed that breast cancer cell lines poorly recaptured mutation patterns of metastatic breast cancer samples, while their CNV profiles were more consistent. Moreover, it is worth noting that cell lines carried many specific genomic alternations, possibly due to culture effects. Examples included the 25 genes presenting cell-line-specific hypermutation. Such large genetic differences and variations revealed by the comparison indicates the importance of selecting cell lines to represent heterogeneous metastatic cancer samples. This study investigated two important factors (i.e., metastatic site and cancer subtype) that need to be considered during cell line selection.

Metastatic sites have their distinct microenvironment that has large impact in shaping the genomic profiles of metastatic cancer cells. However, the metastatic-site-specific TC analysis did not identify cell lines with metastatic-site-specific suitability, which seems not to reflect the impact of microenvironment. Further differential gene expression analysis revealed higher expression of immune related genes in metastatic cancer samples (compared to cell lines), suggesting that the media used to culture cancer cell lines did not model tumor microenvironment appropriately. Therefore, we conclude cell lines do not carry indicative genomic signatures that are shaped by the microenvironment of individual metastatic sites and that is the reason why we did not find metastatic-site-specific cell lines.

Breast cancer is quite heterogeneous and we showed that PAM50 subtypes were maintained in metastatic breast cancer cells. Considering the large genomic difference between Basal-like and other subtypes, it is not surprising that in subtype-specific TC analysis Basal-like subtype showed lower correlation with others. Prior to this study, significant research had been completed to select representative cell lines as models for breast cancer, but the subtype information was not taken into consideration. Our analysis reveals the importance and necessity of subtype-specific cell line selection. In the future as data continues to accumulate, more factors can be considered for appropriate cell line selection and we can start building an ad-hoc mapping algorithm: inputs would be the characteristics of metastatic cancer samples (subtype, metastatic site, even age, race, stage etc.) as well as the specific scientific question of interest and the output would be a list of appropriate cell lines.

We picked out suitable cell lines according to subtype-specific TC analysis results. Surprisingly, we found MDA-MB-231, the widely-used triple-negative cell line in metastatic breast cancer research, was dramatically different from Basal-like metastatic breast cancer samples. As Basal-like breast cancer is itself highly heterogeneous, we could not exclude the possibility that MDA-MB-231 represents a rare subtype not included in the MET500 dataset. According to our analysis, HCC70 seems to be a better model, but this does not mean it can be directly employed to study cancer metastasis as many other criteria are needed for the assessment.

Organoids are recently established using 3D culture methods. Our analysis suggested that compared to cell lines, they resemble the transcriptome of patient samples more closely, which is a critical characteristic in drug testing. It is also important to note that cell lines evaluated in our study were established much earlier than organoids. During culturing process, they could have accumulated additional genomic alternations, which may partially explain why organoids are more tightly correlated with patient samples. In addition, through gene set differential activity analysis, we showed that some gene sets had organoid-specific activity. Recent studies have shown that organoids preserve the histological architecture, gene expression, and genomic landscape of the original tumor^21^. Together with our comparative studies, we conclude that the value of organoids in translational research remains unknown, while their high genomic similarity with patient samples warrants further investigation.

In summary, by leveraging publicly available genomic data, we comprehensively evaluated the suitability of breast cancer cell lines as models for metastatic breast cancer. Our study introduces a simple framework for cell line selection which can be easily extended to other cancer types. Although there are concerns about data quality and discrepancies between different studies/platforms, our large-scale analysis and cross-platform validation hopefully addresses these concerns and demonstrates the power of leveraging open data to gain biological insights of cancer metastasis. We hope that the recommendations in this study may facilitate improved precision in selecting relevant and suitable cell lines for modeling in metastatic breast cancer research, which may accelerate the translational research.

## Methods

### Datasets

The raw RNA-Seq data of MET500 samples were downloaded from dbGap (under accession number phs000673.v2.p1) and further processed using RSEM^22,23^. FPKM values were used as gene expression measure. To keep consistent with other RNA-Seq datasets, only the RNA-Seq samples profiled with PolyA protocol were considered. The somatic mutation and copy number variation (CNV) data of MET500 samples were downloaded from MET500 web portal (https://met500.path.med.umich.edu/downloadMet500DataSets).

All CCLE data (including gene expression profiled by RNA-Seq and microarray, somatic mutation call and CNV) were downloaded from the CCLE data portal (https://portals.broadinstitute.org/ccle).

For TCGA breast cancer samples, somatic mutation calling results of were downloaded from cBioPortal^24,25^ and RSEM-processed gene expression data were downloaded from UCSC Xena data portal (https://xena.ucsc.edu/)^26^.

The RNA-Seq data of patient-derived organoids was from Biobank^27^.

We also searched GEO and manually assembled another microarray dataset containing gene expression value of 103 metastatic breast cancer samples^28,29,30,31^. The GEO accession numbers used were GSE11078, GSE14017, GSE14018, and GSE54323.

The gene expression data of lung-metastasis-derived MDA-MB-231 were downloaded from GEO under accession number GSE2603.

Detailed statistics of the above datasets are listed in Table S6.

### Identification of differentially mutated genes between MET500 and TCGA samples

Given a gene, we computed the right-tailed p-value to test whether it has significantly higher mutation frequency in metastatic breast cancer samples as follows:

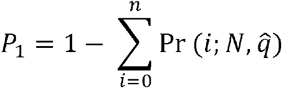

Where Pr is the probability mass function of binomial distribution, N is the number of genotyped MET500 breast cancer cohorts, n is the number of MET500 breast cancer cohorts in which the gene is mutated and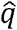,. is the mutation frequency of the gene in TCGA dataset (for genes with zero mutation frequency, we used the minimum mutation frequency across all genes). Similarly, we computed left-tailed p-value (1-P_1_) to test whether a gene has significantly lower mutation frequency in metastatic breast cancer samples. To control FDR, we applied the Benjamini-Hochberg procedure on left-tailed and right-tailed p-values respectively^32^.

### TC analysis with RNA-Seq and microarray data

To perform TC analysis with RNA-Seq data, we first rank-transformed gene RPKM values for each CCLE cell line and then ranked all the genes according to their rank variation across all CCLE cell lines. The 1,000 most-varied genes were kept as “marker genes” (we tried different gene sizes in the early preliminary analysis and did not find the large variation of results, so we decided to choose 1,000 most-varied genes in this study). Given RNA-Seq profiles of a cell line (or an organoid) and several patient samples, we compute spearman rank correlation (across the 1,000 marker genes) between the cell line (organoid) and each sample and the median value of computed spearman rank correlation values was defined as the transcriptome-similarity of the cell line (organoid) with the patient samples. For microarray data, a similar procedure was applied and the 1,000 most-varied probe sets were used to compute correlation values.

We also extended the above method to compute CNV similarity. Instead of selecting “marker genes”, all of the 1,630 commonly genotyped genes were used.

### PAM50 sub-typing and t-SNE visualization

The genefu package was used to determine breast cancer subtype^33,34^. To visualize tumor samples with t-SNE, we first computed the pair-wise distance between every two samples as 1 minus the spearman rank correlation across PAM50 genes and then applied the function Rtsne to perform 2D dimensional reduction^35^.

### PubMed search

The number of PubMed abstracts or full texts mentioning a CCLE breast cancer cell line was determined using the PubMed Search feature on May 10, 2018 (https://www.ncbi.nlm.nih.gov/pubmed/). For each cell line, we searched with a keyword “[cell line name] metastasis”. We repeated this step for the terms “metastatic”, “breast cancer”, and “metastatic breast”. These searches returned highly correlated results, so we used the search terms which returned the most results: “[cell line name] metastasis”.

### Identification of differentially expressed genes and differentially activated gene sets

DESeq2 was used to identify differentially expressed genes (FDR < 0.001 and abs(log2FC) > 1) and DAVID bioinformatics sever was used to perform Gene Ontology enrichment analysis^36,37^. To increase statistical power, only protein coding genes were considered. The 50 hallmark gene sets were downloaded from MSigDB (http://software.broadinstitute.org/gsea/msigdb/) and the R package GSVA was used to perform ssGSEA analysis^38–41^. In DA analysis, for each gene set Wilcoxon rank test was used to assign p-value in the comparison of ssGSEA scores of different subtypes.

### Software tools and statistical methods

All of the analysis was conducted in R and the code is freely available at https://github.com/Bin-Chen-Lab/MetaBreaCellLine. The ggplot2 and ComplexHeatmap packages were used for data visualization^42,43^. Tumor purity was estimated using ESTIMATE^16^. CNTools was used to map the segmented CNV data to genes^44^. If not specified, the Wilcoxon rank test was used to compute p-value in hypothesis testing.

## Acknowledgments

We thank Dr.Nijman of Utrecht Medical Center for providing us organoid RNA-Seq data. The research is supported by R21 TR001743 and K01 ES028047 and the MSU Global Impact Initiative. The content is solely the responsibility of the authors and does not necessarily represent the official views of sponsors.

## Supplementary Figures

**Fig S1.**
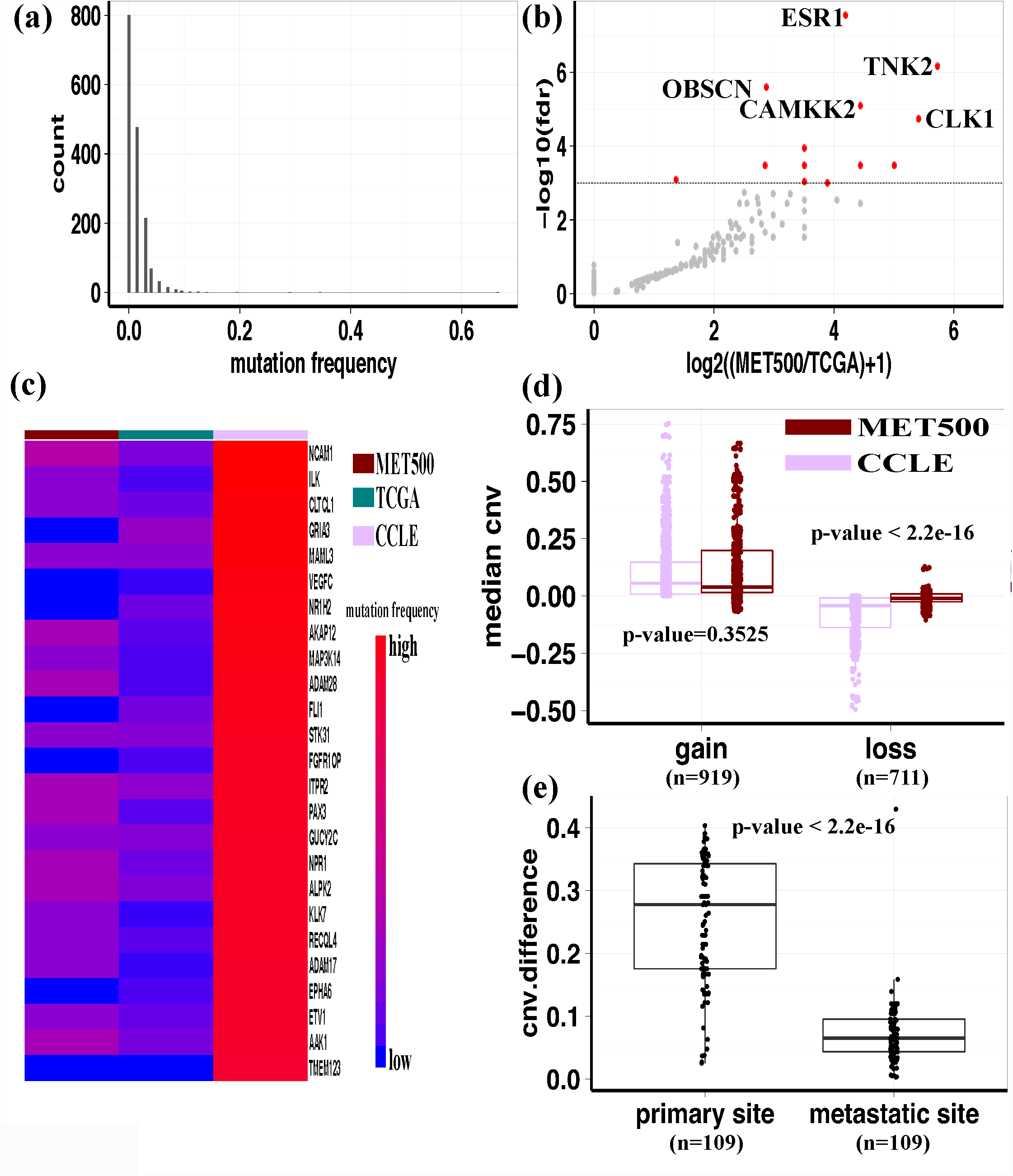
(a) Long-tailed gene mutation spectrum in MET500 breast cancer samples. (b) Volcano plot of differential gene mutation analysis. The dashed line corresponds to FDR = 0.001. (c) Visualization of log10 transformed mutation frequency of the 25 genes that are specifically hyper-mutated in CCLE breast cancer cell lines. (d) Boxplot of median CNV of grouped genes in MET500 breast cancer samples and CCLE breast cancer cell lines. Genes are grouped according to whether showing gain or loss of copy number in CCLE breast cancer cell lines. (e) CCLE breast cancer cell lines derived from metastatic sites more closely resemble the CNV status of genes with high copy-number-gain in MET500 breast cancer samples. Left: absolute value of median-CNV difference between MET500 breast cancer samples and CCLE breast cancer cell lines derived from primary sites. Right: absolute value of median-CNV difference between MET500 breast cancer samples and CCLE breast cancer cell lines derived from metastatic sites.

**Fig S2.**
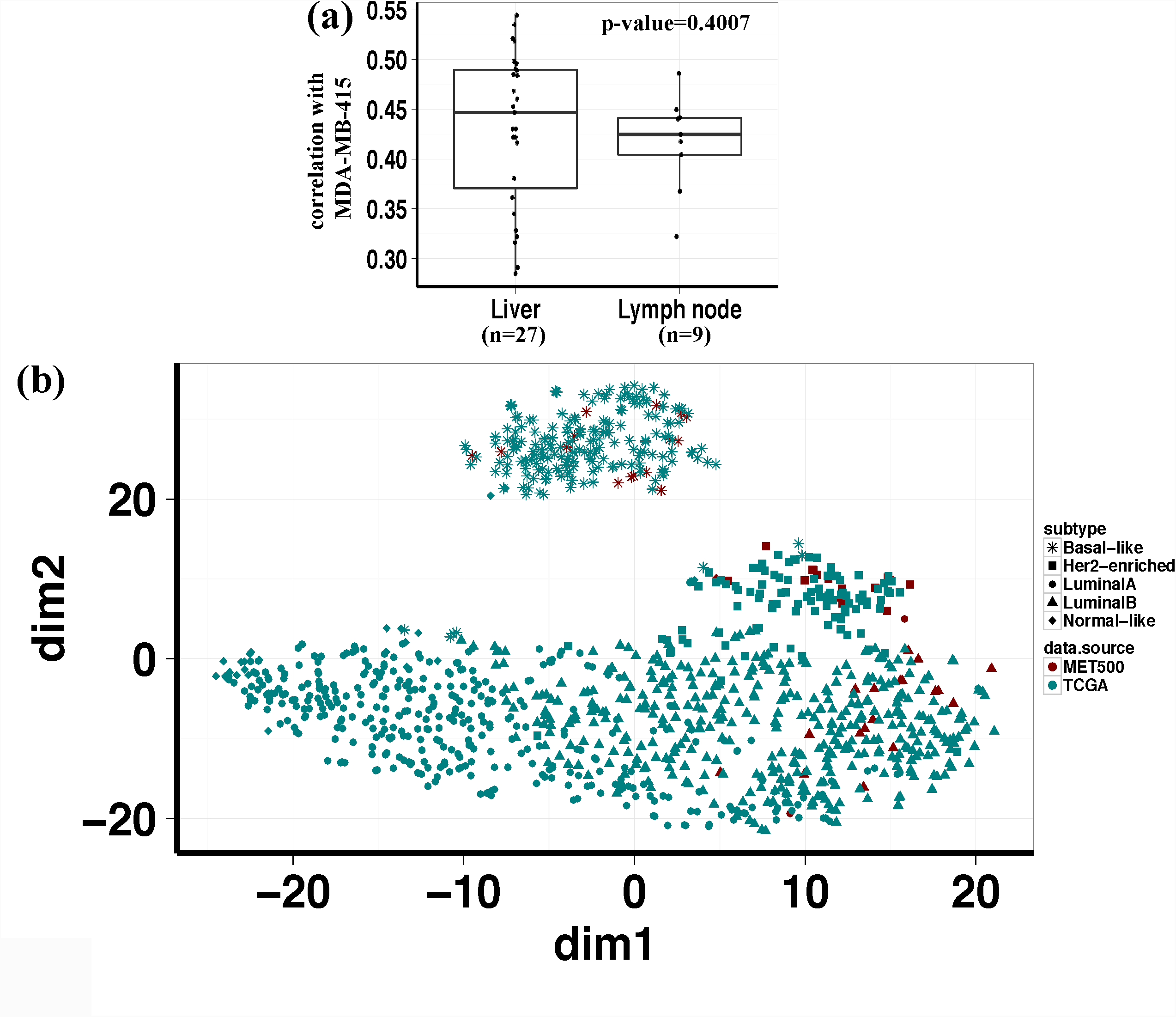
(a) MET500 breast cancer samples derived from liver and lymph node do not show significantly different expression correlation with MDA-MB-415. (b) t-SNE plot of TCGA and MET500 breast cancer samples. Data-sources are labeled by color and subtypes are labeled by shape.

**Fig S3.**
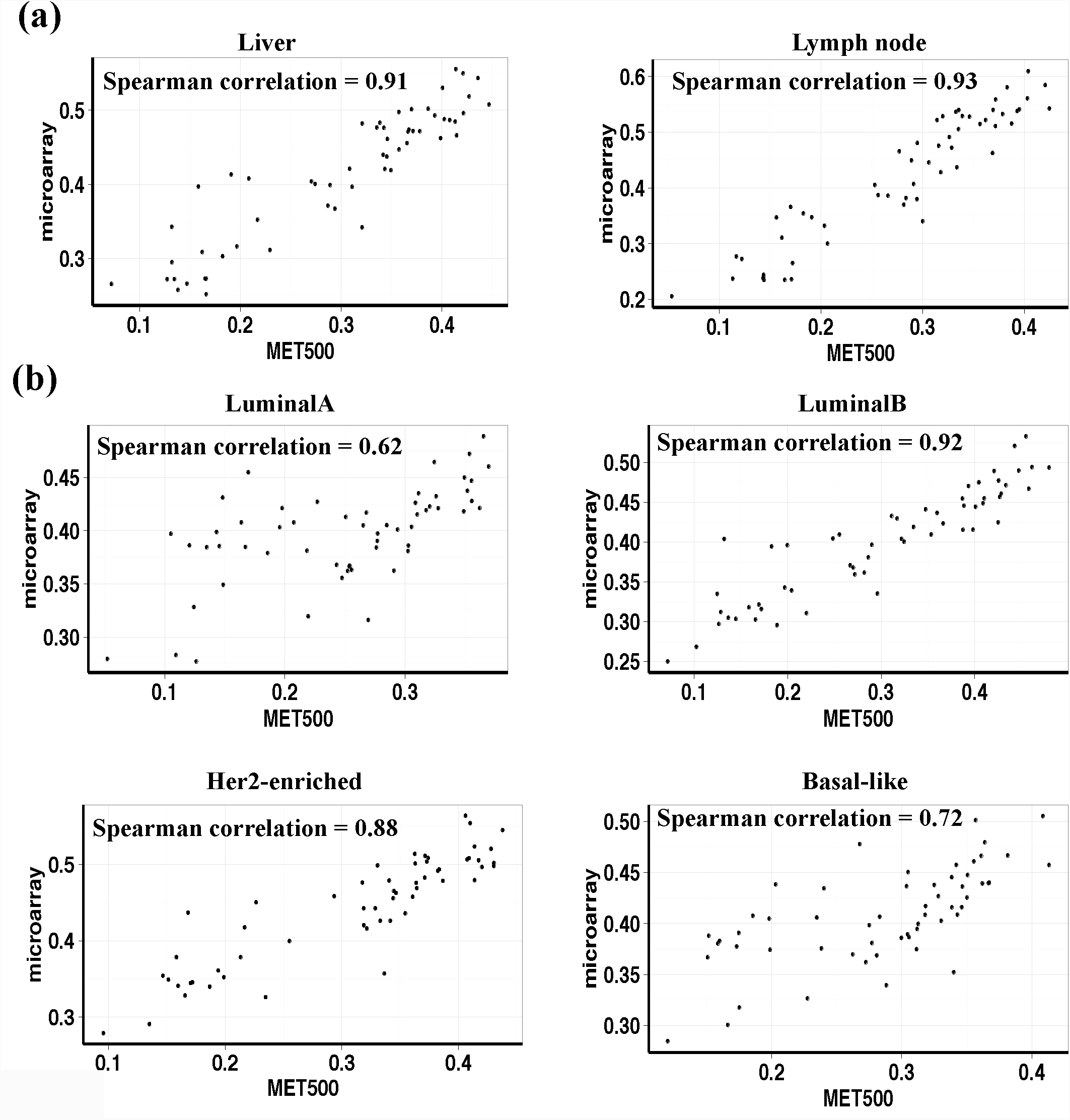
Metastatic-site-specific and subtype-specific TC analysis results are highly correlated between MET500 dataset and the microarray dataset. In each plot, a dot is a CCLE breast cancer cell line, with x-axis representing transcriptome-similarity derived from MET500 dataset, and y-axis representing transcriptome-similarity derived from the microarray dataset.

**Fig S4.**
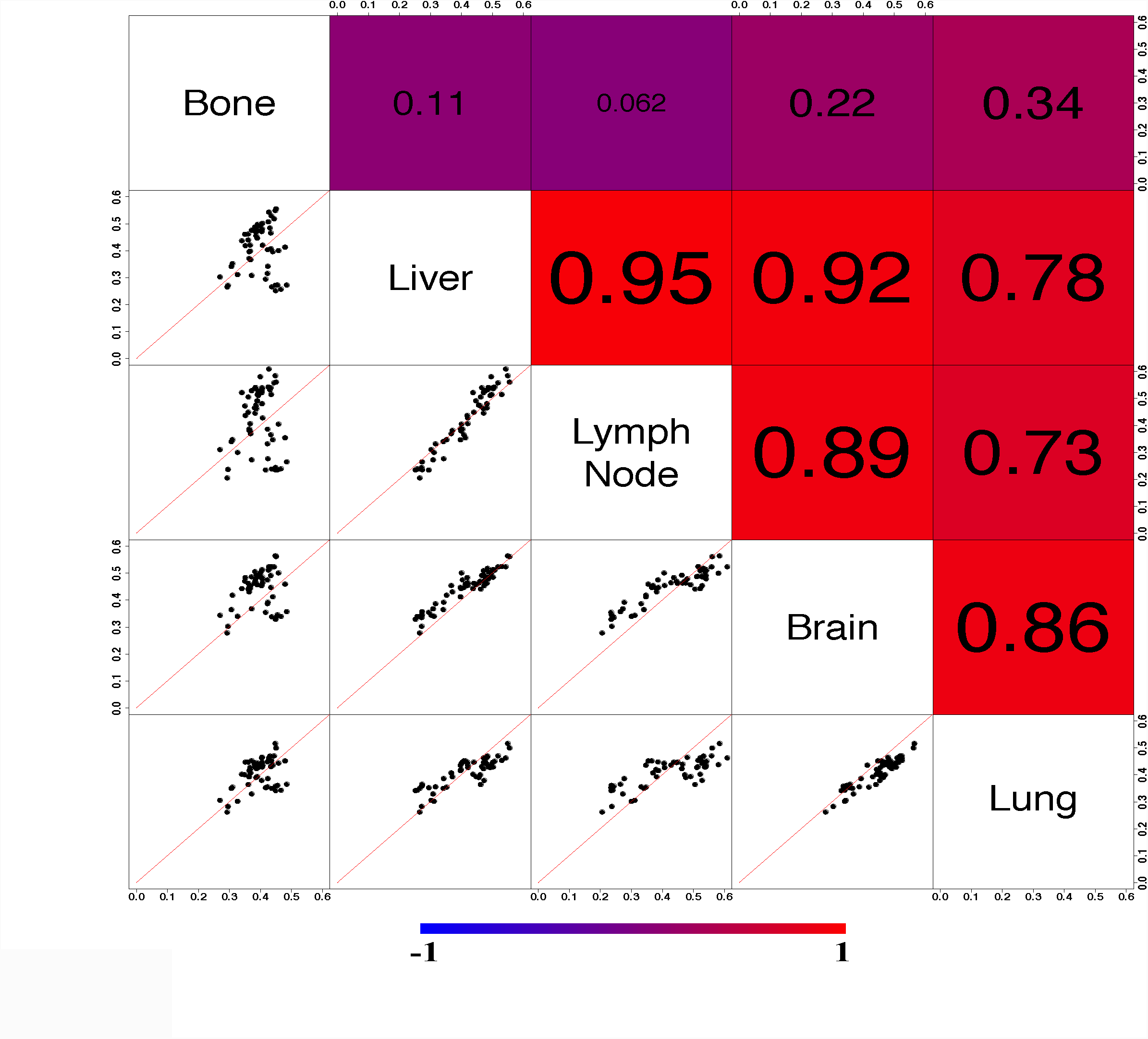
Pair-wise comparison of metastatic-site-specific TC analysis results among metastatic sites (microarray dataset).

**Fig S5.**
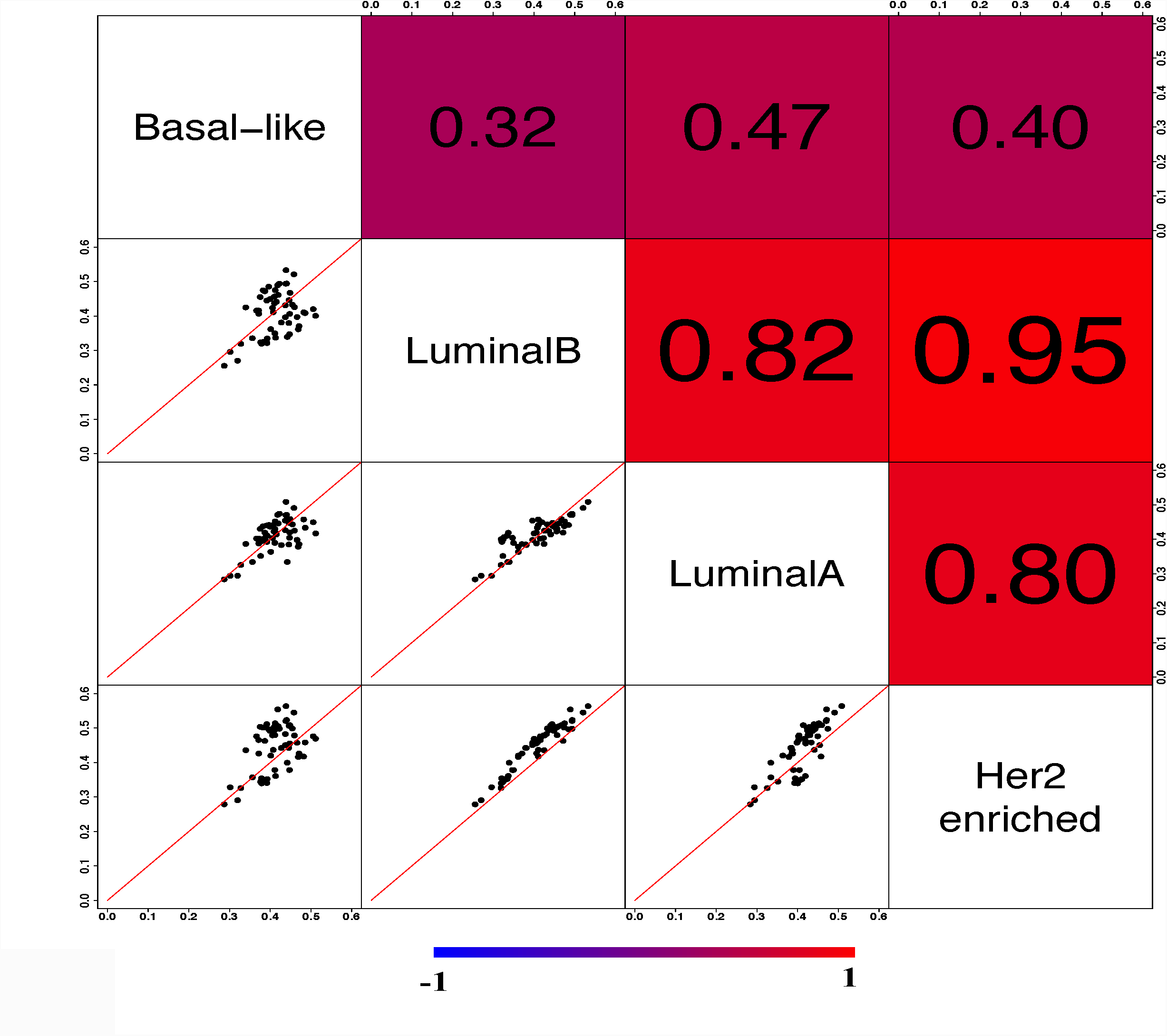
Pair-wise comparison of subtype-specific TC analysis results among subtypes (microarray dataset).

**Fig S6.**
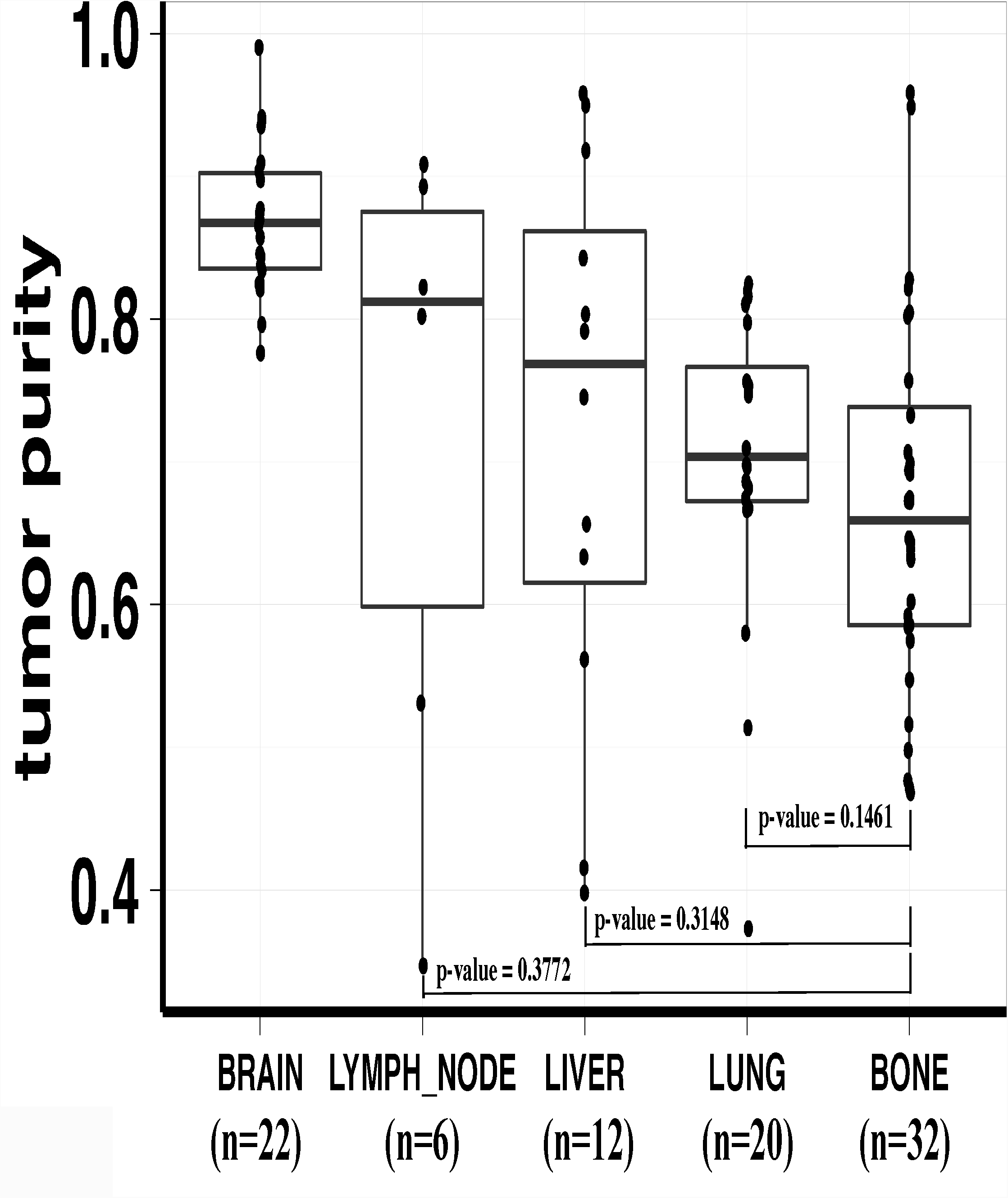
Boxplot of tumor purity (microarray dataset).

**Fig S7.**
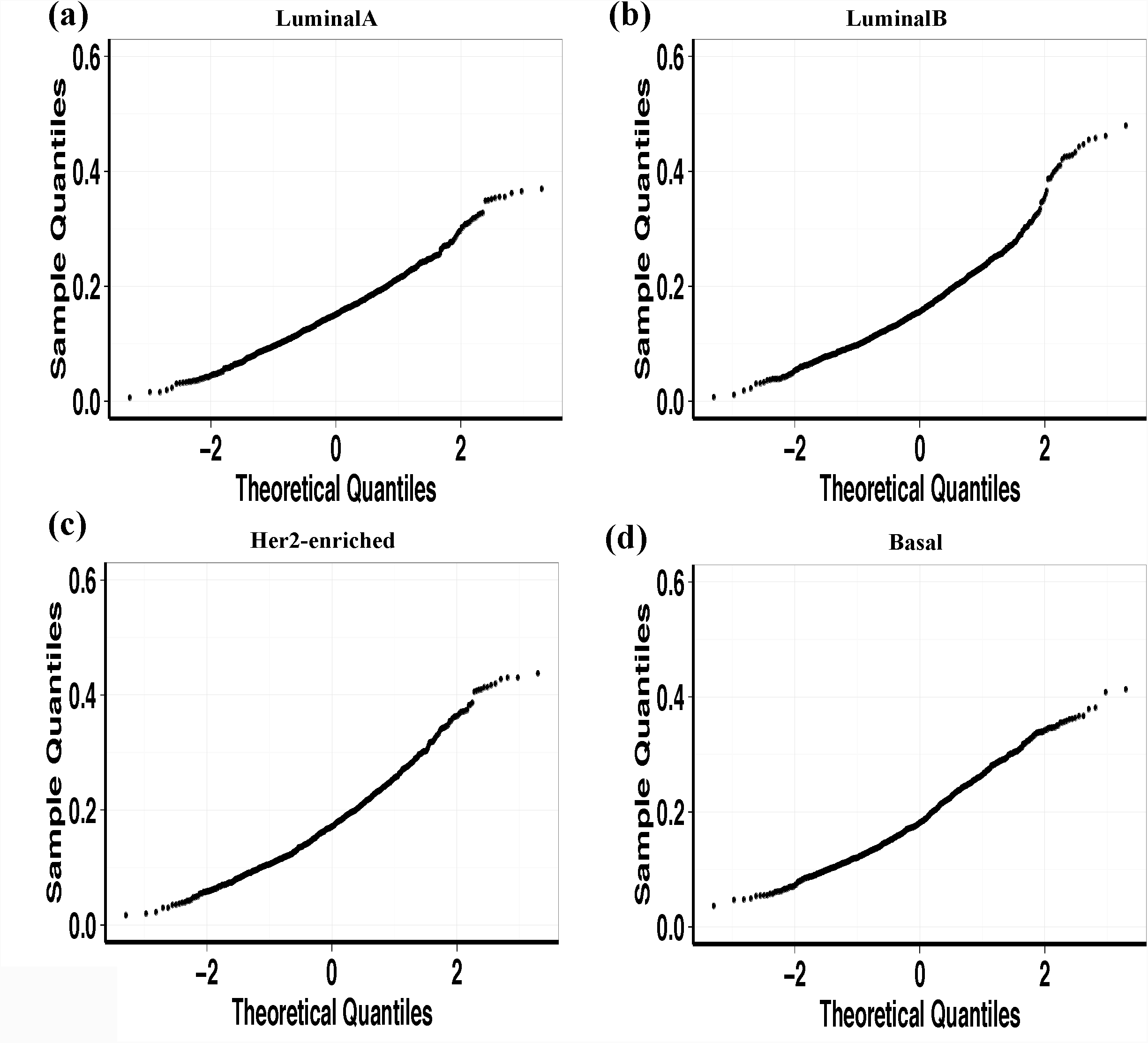
Normal QQ-plot to confirm that the transcriptome-similarity between a random CCLE cell line and MET500 breast cancer samples of a specific subtype approximately follows normal distribution. (a) LuminalA subtype. (b) LuminalB subtype. (c) Her2-enriched subtype. (d) Basal-like subtype.

**Fig S8.**
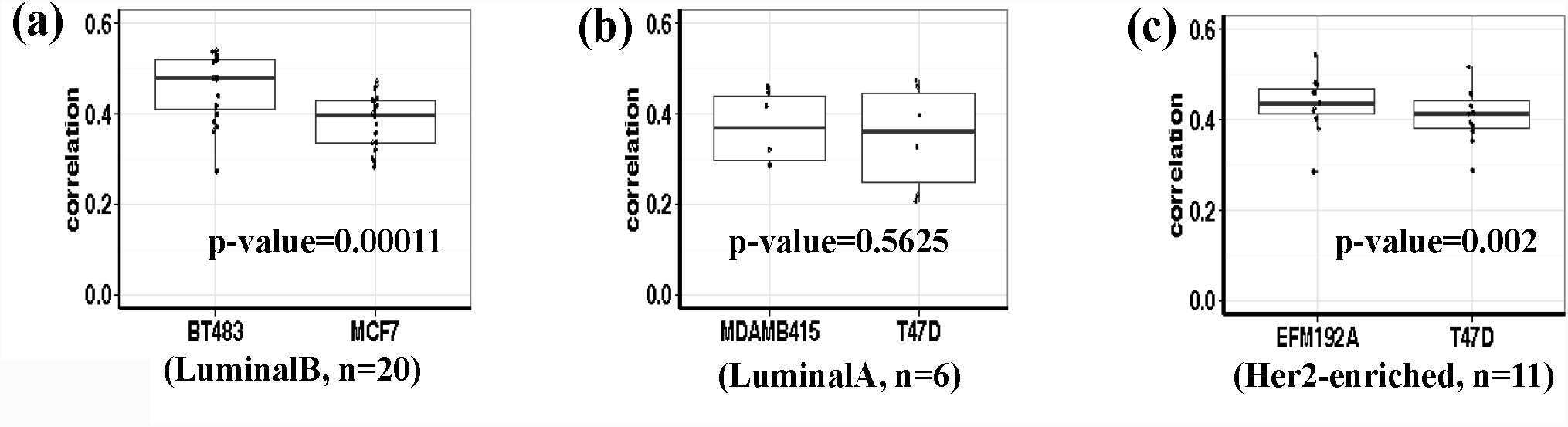
(a) Compared to BT483, MCF7 shows significantly lower expression correlation with LuminalB MET500 breast cancer samples. (b) Compared to MDA-MB-415, T47D does not show significantly lower expression correlation with LuminalA MET500 breast cancer samples. (c) Compared to EFM192A, T47D shows significantly lower expression correlation with Her2-enriched MET500 breast cancer samples.

**Fig S9.**
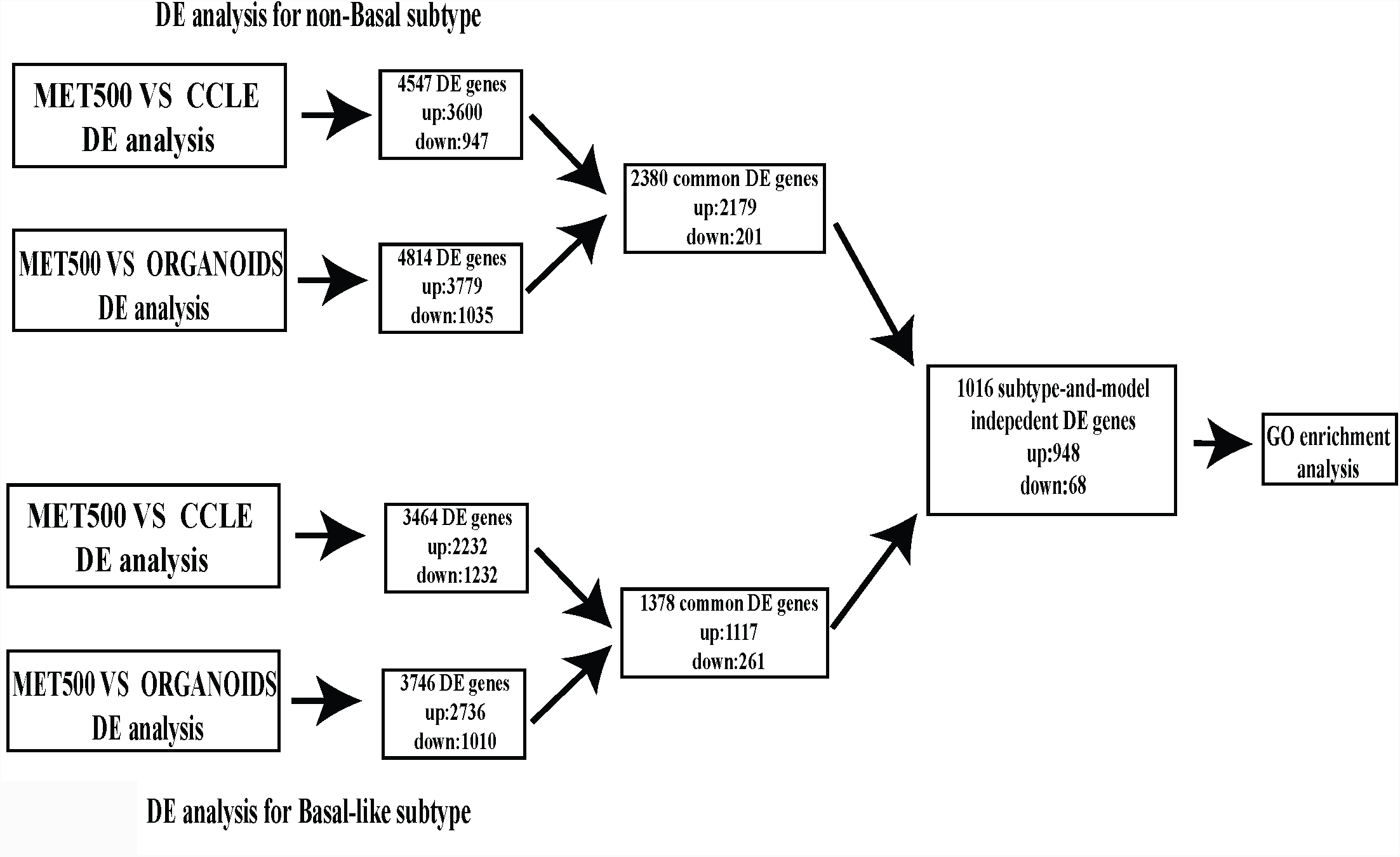
Workflow of differential gene expression analysis.

**Fig S10.**
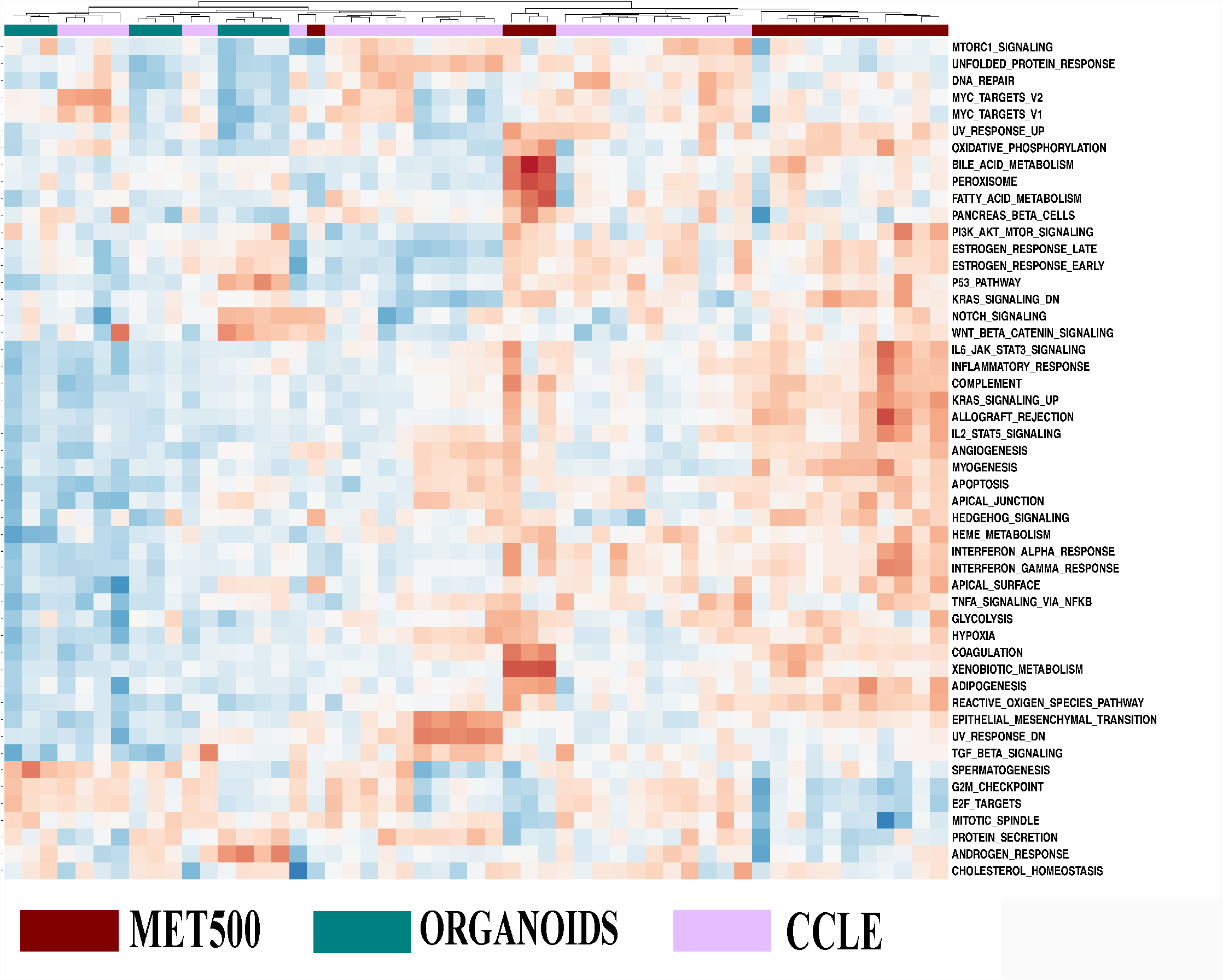
Visualization of ssGSEA scores across Basal-like CCLE breast cancer cell lines, MET500 breast cancer samples and organoids.

## Supplementary Tables

**Table S1.** Mutation frequency of the 75 highly (or differentially) mutated genes in CCLE, TCGA, and MET500 dataset.

**Table S2.** Characteristics of the 57 CCLE breast cancer cell lines.

**Table S3.** Testing suitability of CCLE breast cancer cell lines for different subtypes.

**Table S4.** Results of GO enrichment analysis.

**Table S5.** Results of DA analysis.

**Table S6.** Detailed statistics of the datasets used in our study.

